# Novel Transgenic Humanized Alpha-1 Antitrypsin Deficiency Mouse Model on Murine *SERPINA1* Null Background

**DOI:** 10.1101/2024.10.11.617858

**Authors:** Regina Oshins, Alek M. Aranyos, Skylar Grey, Naweed Mohammad, Yuanqing Lu, Jorge E. Lascano, Tammy Flagg, Karina Serban, Mark Brantly, Nazli Khodayari

## Abstract

Alpha-1 antitrypsin deficiency (AATD) is a rare genetic disorder caused by accumulation of misfolded α-1 antitrypsin within hepatocytes. AATD patients are prone to develop liver disease that remains undiagnosed until the late stages of the disease. Due to challenges in manipulating the α-1 antitrypsin genes in mice, determining a true loss of function of α-1 antitrypsin in previous AATD mouse models has been challenging. Here, we report generation and liver characterization of a new humanized transgenic mouse model for AATD with a background of a CRISPR-Cas9 generated *SERPINA1*-null mouse. Male and female transgenic mice for normal (Pi*M) and mutant (Pi*Z) variants of human α-1 antitrypsin at 4-6 months of age were subjected to this study. The accumulation of human α-1 antitrypsin in the hepatocytes and fibrotic features of the liver were monitored by performing an in vivo study. We demonstrate a strong liver phenotype satisfying clinically relevant manifestations of liver pathology associated with AATD, including hepatic accumulation of human α-1 antitrypsin globules, liver deposition of extracellular matrix proteins, hepatic ER stress, and liver fibrosis in Pi*Z mice, in addition to mild systemic inflammation. In addition to major phenotypic criteria of AATD-associated liver fibrosis, accompanying single-nucleus RNA-seq data demonstrate activation of pathways associated with liver metabolic changes, inflammation, and regeneration. Data from this study suggest our humanized transgenic AATD mouse model could provide a suitable model to study α-1 antitrypsin loss of function, replicate the pathophysiology of AATD associated liver disease, and evaluate therapeutic reagents against this disease.

**NEW & NOTEWORTHY:** We have characterized a new humanized transgenic mouse model of α-1 antitrypsin deficiency with a *SERPINA1*-null background that shows strong manifestations of liver disease. Our data explores the altered phenotype of α-1 antitrypsin deficient hepatocytes and suggests a relationship between liver cell types during disease progression. This model may become a useful tool for investigating α-1 antitrypsin loss of function, pathogenic mechanisms, and for drug discovery aimed at both prevention and treatment of the disease.

## INTRODUCTION

Alpha-1 antitrypsin is the most abundant serine protease inhibitor in the plasma and highly expressed in the liver (1). Alpha-1 antitrypsin deficiency (AATD), a rare genetic disease affecting more than 100,000 individuals in the U.S. is caused by the Z mutation of the *SERPINA1* gene (E342K) which encodes α-1 antitrypsin. The Z mutant variant of α-1 antitrypsin misfolds and aggregates within hepatocytes. Hepatic accumulation of polymerized Z α-1 antitrypsin causes endoplasmic reticulum (ER) stress followed by chronic liver injury ranging from inflammation and fibrosis to cirrhosis and hepatocellular carcinoma (2, 3). Currently, there is no treatment for AATD-associated liver disease except for liver transplantation (4).

An animal model capable of consistently replicating liver disease associated with AATD would be a valuable tool for understanding the pathophysiology and disease mechanisms as well as for *in vivo* evaluation of therapeutic agents to enhance progress of potential drug development. A number of different animal models have been previously developed and are currently used to study AATD-associated liver disease (5, 6). While these models are beneficial, it is important to note that these models continue to express endogenous mouse α-1 antitrypsin protein. This aspect could be a limitation when studying α-1 antitrypsin loss of function (7), especially in the liver, where the pathology was always attributed to α-1 antitrypsin gain-of-function and it might not replicate the liver disease associated with human AATD, impacting translatability of the results (8). Additionally, the presence of endogenous mouse α-1 antitrypsin protein hinders the examination of the anti-inflammatory and immunomodulatory functions of α-1 antitrypsin (9), posing another challenge for the clinical application of the study’s findings.

In order to address the limitations of previous animal models of AATD, we have developed a new transgenic mouse line expressing human wildtype (Pi*M) or Z variant α-1 antitrypsin (Pi*Z) from murine Serpina1-null background mice (10) to study of AATD-associated liver disease. By microinjection insertion of either the human M or Z variant of the *SERPINA1* gene which includes both the macrophage and hepatocyte specific promoter into the oocytes of C57BL/6J-Serpina1 knockout mice (11), we were able to achieve appropriate α-1 antitrypsin RNA and protein expression in the liver of mice which allows for study of AATD-associated liver pathologies. In this study, we assessed hepatic accumulation of misfolded α-1 antitrypsin by immunohistochemistry (IHC) analysis and have validated the results with a series of techniques, including western blotting and circulating plasma levels of human α-1 antitrypsin in the transgenic mice were measured using nephelometry. We show this ATTD mouse model displays mild systemic inflammation and stable liver manifestations of the disease covering abundant AATD pathological features such as hepatic accumulation of α-1 antitrypsin globules and a significant loss of plasma circulating α-1 antitrypsin. This newly designed AATD mouse line represents a novel tool to study and elucidate underlying mechanisms of AATD-associated tissue injuries and is suitable for the assessment of response to therapy with drugs known to be efficacious against AATD.

## MATERIALS AND METHOD

### Transgenic AATD Mouse Model and Treatments

All mice listed were bred with approval from the University of Florida Institutional Animal Care and Use Committee (IACUC no. 202200000740) with adherence to the Guide for Care and Use of Laboratory Animals (NIH, 86-23). The oocytes from female mice homozygous for the *Serpina1* knockout ( C57BL/6J-*Serpina1^em3Chmu^*/J knockout, The Jackson Laboratory, Strain#:035015) were harvested and microinjected by The Jackson Laboratory with human wildtype (Pi*M) or mutant (Pi*Z) variant of α-1 antitrypsin full-length DNA construct (Genbank ID: NC_000014.9; sequence 694374747 to 94395692) which contains both the macrophage and hepatocyte enhancer-promoter regions. Embryos were transferred into pseudo pregnant females, and the resulting pups were genotyped for the presence of human *Serpina1* transgenes. Correctly, targeted pups were then bred to the background strain (C57BL/6J-*Serpina1^em3Chmu^*/J, homozygous for *Serpina1* knockout) for germline transmission. Transgenic Pi*M and Pi*Z mice hemizygous for the human transgenes and their littermates (C57BL/6J-*Serpina1^em3Chmu^*/J) were bred and maintained under pathogen-free conditions at 22 ± 3°C with a 12:12 h light-dark cycle and fed normal rodent chow (12) within our in-house breeding colony under guidance from IACUC. Prior to using animals from these colonies, genotype is checked utilizing an allelic discrimination assay (TaqMan SNP Genotyping). Phenotype is confirmed by blood draw to test plasma levels of human α-1 antitrypsin by nephelometry. Mice (n=6, 4 males, 2 females) were 4-6 months old at time of sacrifice and sex was not accounted for in the study due to lack of sex differences observed in AATD. To induce systemic inflammation, Mice (n=6, 3 males, 3 females) were injected intraperitoneally (i.p.) with LPS (3 mg/kg) and followed for 24 h. LPS (*Escherichia coli* 0111: B4, Sigma-Aldrich, Milwaukee, WI, USA) was freshly dissolved in sterile phosphate buffered saline (PBS) (13).

### Quantitative Real-Time PCR

RNA isolation and cDNA synthesis were performed as previously described (12). Briefly, RNA from liver tissues was extracted using a RNeasy Plus Mini Kit and reverse transcribed using Superscript VILO reverse transcriptase. PCR was performed using TaqMan Fast Advanced Master Mix on a 7500 real-time PCR system (Applied Biosystems). All probes were purchased from ThermoFisher (Carlsbad, CA). Gene expression levels were normalized to levels of ribosomal 18S housekeeping gene. Data are expressed as fold change over control and were calculated using either the 2^−ΔC(T)^ or 2^−ΔΔC(T)^ method.

### Tissue Preparations and Immunohistochemistry

Pi*Z and Pi*M livers were formalin-fixed paraffin-embedded (FFPE) or embedded in Optimal Cutting Temperature (OCT) after 4% paraformaldehyde fixation and 30% sucrose cryopreservation. Tissues were then sent to the University of Florida’s Molecular Pathology Core (RRID:SCR_016601) for sectioning and staining.

### Conventional Transmission Electron Microscopy Processing

Perfused Pi*M (n=1) and Pi*Z (n=1) liver tissues were trimmed and fixed in 4% paraformaldehyde and 2.5% glutaraldehyde in 0.1M sodium cacodylate buffer containing 2mM MgCl_2_, 1mM CaCl_2_, 0.25% NaCl at pH 7.42. All subsequent steps were microwave-assisted, processed with a Pelco BioWave Pro laboratory microwave (Ted Pella, Redding, CA, USA) and an SBT digital orbital shaker (Southwest Science, Trenton, NJ, USA). Tissues pieces were buffer washed, post-fixed in buffered 2% osmium tetroxide, water washed, and dehydrated in a graded ethanol series 25% to 100% in 5-25% increments, followed by 100% anhydrous acetone. Dehydrated samples were infiltrated in graded acetone-Araldite/Embed epoxy resin with Z6040 embedding primer (Electron Microscopy Sciences, Hatfield, PA) at 30%, 50%, 70%, and 100% and cured at 70°C. Semi-thick sections (500 nm) were collected and stained with toluidine blue. Ultra-thin sections (120 nm) were collected on a carbon-coated Formvar 100 mesh copper grid (EMS, Hatfield, PA), and post-stained with 2% aqueous uranyl acetate and lead citrate.

### Transmission Electron Microscopy Imaging

Sections were either examined with a FEI Tecnai G2 Spirit Twin TEM (FEI Corp., Hillsboro, OR) operated at 120 kV and digital images were acquired with a Gatan UltraScan 2k × 2k camera and Digital Micrograph software (Gatan Inc., Pleasanton, CA) or a Thermo Scientific Talos L120C G2 TEM and digital images were acquired with a Thermo Scientific Ceta CMOS 4k x 4k camera and Velox software (ver. 3.11).

### Protein Analysis

Protein concentrations of whole liver lysates were quantified by BCA, prepared in denaturing conditions, and analyzed on a 4-20% gradient tris-glycine gel. The gel was transferred to a 0.45µm nitrocellulose membrane (Bio-Rad, Hercules, CA). After blocking, membranes were incubated with primary antibody overnight at 4°C. GRP75, VDAC1, HSP60, ATF4, and GAPDH rabbit anti mouse primary antibodies were obtained from ProteinTech (Rosemont, IL). BiP antibody was purchased from BD Biosciences (Franklin Lakes, NJ) and CHOP antibody was purchased from Cell Signaling (Danvers, MA). Horseradish peroxidase (HRP)-conjugated goat anti-mouse or anti-rabbit IgG antibodies (Bio-Rad) were used as secondary antibodies. Membranes were probed using secondary antibody for one hour and imaged on a Bio-Rad imager (Hercules, CA) using a chemiluminescent substrate.

### Nephelometry

Circulating human α-1 antitrypsin plasma levels of transgenic mice were determined by nephelometry (Behring Diagnostics, Marburg, Germany) using in-house standards and controls.

### Antibody Labeling of Cells and Flow Cytometry

Liver nonparenchymal cells isolated from WT and null mice with and without LPS treatment (*n* = 6/group) were stained as previously described (6) and analyzed using a CytoFlex (Beckman Coulter, Brea, CA). Dead cells were excluded using Live/Dead Near IR (Invitrogen, Carlsbad, CA). Cells were stained with CD45-PerCP, CD11b-BV750, Ly6C-PE/Cy7, and Ly6G-BV605 antibodies from BioLegend (San Jose, CA). Data were analyzed using FlowJo (BD Biosciences, San Jose, CA) software. Our gating strategy was first gating out debris and cell aggregates. Next, live CD45+ liver NPC were identified and the expression of CD11b was confirmed in the population. Ly6C and Ly6G have been further used to identify neutrophil and monocyte populations.

### Single-nucleus RNA Sequencing

Single nucleus RNA sequencing was conducted by Singulomics Corporation (https://singulomics.com/, Bronx NY). In summary, frozen Pi*M (n=1, male) and Pi*Z liver tissues (n=1, male) were homogenized and lysed with Triton X-100 in RNase-free water for nuclei isolation. The isolated nuclei were purified, centrifuged, and resuspended in PBS containing BSA and RNase inhibitor. The isolated nuclei (4000 nuclei/sample) were loaded to 10x Genomics Chromium Controller to encapsulate single nuclei into droplet emulsions following manufacturer’s recommendations (10x Genomics, Pleasanton, CA). Library preparation was performed according to instructions in the Chromium Next GEM 3’ Single Cell Reagent kits v3.1. Amplified cDNAs and libraries were measured by Qubit dsDNA HS assay (ThermoFisher Scientific, Wilmington, DE) and quality was assessed by TapeStation (Agilent Technologies, Santa Clara, CA). Libraries were sequenced on a NovaSeq instrument (Illumina, San Diego, CA), and reads were subsequently processed using 10x Genomics Cell Ranger analytical pipeline (v7.0.1) with the mouse reference genome mm10 with introns included in the analysis.

### Differentially Expressed Gene Analysis

Barcode matrices containing single nucleus RNA sequencing data were read into R (v 4.3.2) and formatted into Seurat objects by the R package Seurat version 5 (14–16) for each of the two samples separately. The two objects were merged, log-normalized, and scaled before going through a standard analysis pipeline. Then we integrated the combined object using canonical correlation analysis (CCA) integration (17). We performed shared nearest neighbor (SNN) clustering using Seurat and then identified the cell types of each cluster using the sc-type package (18). Clusters identified by sc-type as “Unknown” were identified manually. Differentially Expressed Genes (DEGs) were identified by the using the function FindMarkers in Seurat. All p-values were adjusted to q-values, which controls the false discovery rate (FDR) using the Benjamini-Hochberg FDR adjustment. Only those genes with both q-value less than 0.05 and log-2-fold change > |1.5| were included. An exception was made for Kupffer cells because there were so few cells; we used p-values rather than q-values for these.

### Functional Enrichment Analysis

To further enhance our understanding of the DEGs, functional enrichment analyses of DEGs were performed using the R package clusterProfiler (19, 20) with default parameters. Gene Ontology (GO) was used to compare DEGs to biological processes and Kyoto Encyclopedia of Genes and Genomes (KEGG) was used to compare DEGs to functional pathways. These analyses helped identify pathways that provide insights into the biological mechanisms driving the observed differences between the experimental groups.

### Statistical Analysis

All results presented in this study are expressed as means ± SD. Statistical analyses were performed using the Prism 9 software program (GraphPad Software) by Student’s t test or Mann–Whitney U test. One-way ANOVA test followed by Newman-Keuls test was performed for multiple-group comparisons, while non-parametric Kruskal-Walli’s test was used when data cannot pass the normality test. Values of P < 0.05 were considered statistically significant.

## RESULTS

### Validation of Human *SERPINA1* Gene and Transgenic Mice

In humans, α-1 antitrypsin is encoded by the *SERPINA1* gene in which a single point mutation (commonly referred to PiZ) causes misfolding and aggregation of misfolded α-1 antitrypsin in hepatocytes resulting in liver injury. Therefore, a 23-kb pMA-pHuman-SerpinA1 construct was used to insert a humanized copy of the *SERPINA1* gene, either expressing the normal M variant of human *SERPINA1* or the Z mutated variant of human α-1 antitrypsin into the mouse genome. The Z point mutation was introduced to the pMA-pHuman-SerpinA1 construct to induce the aggregation propensity of α-1 antitrypsin associated with AATD. The transgenic construct contains the entire human *SERPINA1* (serine peptidase inhibitor, clade A, member 1 cluster) gene under its endogenous promoters (both hepatocyte and macrophage) plus 5 kB of the 5’ and 3 kb of the 3’ flanking genomic DNA sequences; the Z construct also contains a glutamic acid to lysine substitution at residue 366 (E366K, also designated 342; G to A; rs 28929474) (Figure 1A and B).

**Figure 1.** Validation of human *SERPINA1* gene inserted into the background mice. (A) A 23-kb pMA-pHuman-SerpinA1 construct map, either expressing the M or Z variant of human α-1 antitrypsin. (B) The transgenic Pi*M and Pi*Z mice were backcrossed more than 10 generations and maintained on a C57BL/6J-Serpina1 knockout mice background. (C) Plasma levels of human α-1 antitrypsin protein expressed in transgenic Pi*M and Pi*Z mice. (D) The plasma levels of human α-1 antitrypsin protein in Pi*M and Pi*Z mice for 20 weeks (weeks 4-24 of age). p-values have been calculated using student t-test method. Pi*M, normal variant of human α-1 -antitrypsin; Pi*Z, mutant human α-1 -antitrypsin.

The transgenic litters expressing M variant α-1 antitrypsin (Pi*M) and Z variant α-1 antitrypsin (Pi*Z) were backcrossed more than 10 generations and maintained on a C57BL/6J-Serpina1 knockout mouse background. The offspring are viable and fertile, although decreased litter productivity has been observed in Pi*Z transgenic mating pairs. We have confirmed the presence of human α-1 antitrypsin protein in the plasma of Pi*M and Pi*Z mice using nephelometry, and results revealed the expression pattern of transgenic human α-1 antitrypsin closely resembled the pattern of human α-antitrypsin (Figure 1C). We have also closely monitored the plasma levels of human α-1 antitrypsin protein in Pi*M and Pi*Z mice for 20 weeks (4-24 weeks of age) and observed no significant changes in α-1 antitrypsin levels in both groups of mice (Figure 1D).

### Liver Characterization of Wildtype and α-1 Antitrypsin Null Mice

The physiological role of α-1 antitrypsin in the liver remains unexplored due to the presence of endogenous mouse α-1 antitrypsin protein in prior AATD transgenic mice. To assess the role of α-1 antitrypsin in the liver, we injected WT and null mice intraperitoneally (i.p.) with LPS (3 mg/kg) and 24 hours later, measured the plasma of levels of AST and ALT activity, markers of liver injury (21). We noticed plasma levels of AST and ALT were identical in untreated WT and null mice. However, AST and ALT activity were significantly higher in the plasma of null mice after LPS treatment, indicating higher levels of liver injury induced by systemic inflammation in null mice as compared to WT mice in response to LPS (Figure 2A). To characterize the immune cell populations in the liver of WT and null mice, we assessed the phenotype of neutrophils and monocyte populations in WT and null (*n* = 6/group) livers using flow cytometry. Nonparenchymal liver single-cell suspensions were first gated on single cells followed by live cells. Neutrophils were identified by expression of CD45, CD11b and Ly6G and lack of Ly6C. Patrolling monocytes were characterized by expression of CD45, CD11b, low expression of Ly6C and lack of Ly6G. Inflammatory monocytes were identified by expression of CD45, CD11b, high expression of Ly6C and lack of Ly6G (Figure 2B). Although both WT and null livers contained the same number of neutrophils (Figure 2C) and patrolling monocytes (Figure 2D), null livers displayed higher levels of Ly6C high cells (inflammatory monocytes) (Figure 2E). Furthermore, we characterized the immune cell populations in the liver of Pi*M and Pi*Z mice, using flow cytometry and compared the results with WT and null mice (*n* = 6/group) as described above. The results indicated that expressing the normal variant of human α-1 antitrypsin in the livers of Pi*M mice reverts the null liver inflammatory phenotype. However, expressing mutant variant of human α-1 antitrypsin failed to revert the null liver inflammatory phenotype in Pi*Z mice (Figure 2E).

**Figure 2.** Liver function and inflammatory response to LPS in *SERPINA1*-null as compared with wildtype mice. (A) The levels of AST and ALT activity in the plasma of both WT and null mice indicated no difference between the liver function of control WT and null mice. However, AST and ALT activity are found to be significantly higher in the plasma of null mice as compared to WT mice in response to LPS induced inflammation. (B) Nonparenchymal cells phenotype (CD45^+^) in control and LPS treated WT (*n* = 6/group) and null (*n* = 6/group) livers performing flow cytometry. CD45 expressing cells are considered as nonparenchymal cells. Neutrophils were analyzed for presence of CD11b and Ly6G and absence of Ly6C surface markers. Ly6C^low^-expressing monocytes are considered as patrolling monocytes and Ly6C^high^-expressing monocytes are considered as inflammatory monocytes. (C) Histogram of neutrophil population, (D) patrolling monocytes, and (E) inflammatory monocytes of the livers using flowcytometry assay in control WT, null, Pi*M, and Pi*Z mice (*n* = 6/group). p-values have been calculated using 2-way ANOVA method. LPS; lipopolysaccharide, Pi*M, normal variant of human α-1 -antitrypsin; Pi*Z, mutant human α-1 -antitrypsin.

### Histopathology of Liver Tissues of Transgenic Mice

Mouse liver tissues stained with hematoxylin-eosin (H&E) examined by light microscopy show normal hepatocytes and sinusoids in both Pi*M and Pi*Z mice compared to wildtype and null background mice. Portal areas could be identified and the presence of portal areas along with central veins showed a lobular structure of the mouse liver tissue (Figure 3A). While we did not observe α-1 antitrypsin accumulation in the liver of wildtype, null, or Pi*M mice, Pi*Z mice exhibited accumulation of Periodic Acid Schiff globules after treatment with Diastase (PASD), indicating α-1 antitrypsin containing globules, at 4 months of age (Figure 3B). To assess liver fibrogenesis associated with AATD, Picro-Sirius Red staining was performed on liver sections from all mouse groups and the results revealed formation of collagen fibers and excessive collagen deposition in the liver of Pi*Z mice compared to background and Pi*M mice (Figures 3C and E). This finding was confirmed by expression of α-smooth muscle actin (α-SMA) in the liver of Pi*Z mice observed by immunohistochemical staining (Figures 3D and F), which indicates presence of activated hepatic stellate cells (HSC) and liver fibrogenesis in the liver of Pi*Z mice associated with hepatic accumulation of α-1 antitrypsin.

**Figure 3.** Liver histopathology of transgenic Pi*M and Pi*Z mice compared to wildtype and *SERPINA1*-null background. (A) Hematoxylin and Eosin staining indicating normal hepatocytes and sinusoids in both Pi*M and Pi*Z mice liver tissues compared to wildtype and null background mice. (B) α-1 antitrypsin accumulation is not detectable in the liver of wildtype, null, and Pi*M mice. Pi*Z mice exhibited accumulation of Periodic Acid Schiff after treatment with Diastase (PASD) staining (dark pink globules) at 4 months of age indicating accumulation of α-1 antitrypsin globules. (C) Pico Sirius Red staining on the liver sections from wildtype, null, Pi*M, and Pi*Z mice revealing formation of collagen fibers and collagen deposition in the liver of Pi*Z mice compared to background mice and Pi*M. (D) Expression of α-SMA (brown signal) in the liver of Pi*Z mice compared to wildtype, null, and Pi*M mice assessed by immunohistochemical staining which is increased in the liver of Pi*Z mice. All images have been captured in ×10 magnification (*n* = 4). p-values have been calculated using student t-test method. Pi*M, normal variant of human α-1 -antitrypsin; Pi*Z, mutant human α-1 -antitrypsin.

### Plasma Markers and Inflammatory Mediators of Transgenic Mice

The hepatic accumulation of Z variant α-1 antitrypsin has been linked to hepatotoxicity, highlighting its role in AATD-associated liver pathology and liver disease progression (22). Therefore, we assessed hepatotoxicity in the plasma of our transgenic mouse model by measuring serum levels of AST and ALT activity. Consistent with previous findings regarding AATD-associated liver disease (23), our Pi*M and Pi*Z mice show no difference between plasma AST activity levels from WT and null mice (Figure 4A). However, ALT activity was significantly higher in the plasma of Pi*Z mice as compared to Pi*M, as well as WT and null mice (Figure 4B). Furthermore, we analyzed circulating inflammatory mediators in the plasma of Pi*M and Pi*Z mice to assess the effect of liver phenotype on systemic inflammation. While there was no difference between plasma cytokines of the background control animals (data not shown) and Pi*M mice, there were increased levels of some inflammatory mediators such as IL-2 and IL-18 in the plasma of Pi*Z compared to Pi*M mice (Figure 4C).

**Figure 4.** Liver enzymes activity and inflammatory mediators of the plasma from transgenic Pi*M and Pi*Z mice compared to wildtype and *SERPINA1*-null background. (A) The levels of AST activity in the plasma of both Pi*M and Pi*Z mice indicated no difference between the liver function of transgenic mice and WT and null mice. (B) ALT activity is found to be significantly higher in the plasma of Pi*Z mice as compared to Pi*M, as well as WT and null mice. (C) The levels of cytokines in the plasma of Pi*M and Pi*Z mice as measured by ELISA, showing significant increase in the levels of IL-2 and IL-18 in the plasma of Pi*Z as compared to Pi*M mice. p-values have been calculated using 2-way ANOVA method. Pi*M, normal variant of human α-1 -antitrypsin; Pi*Z, mutant human α-1 -antitrypsin.

### Ultrastructural Characterization of the AATD Liver Tissue

To investigate the AATD hepatic phenotype in our transgenic mouse model in more detail, we performed transmission electron microscopy (TEM). TEM investigation of liver sections from a Pi*M mouse revealed normal ultrastructure. The rounded nuclei contained normal chromatin, pores, and membranes. The hepatocytes contained numerous round and ovoid mitochondria with well-developed cristae and were seen scattered throughout the cytoplasm. A well-developed rough ER around the mitochondria in the cytoplasm could also be seen, studded with numerous ribosomes (Figure 5A). However, Pi*Z hepatocytes contained nuclei with irregular membranes, lipid droplets, and abnormal mitochondria. We observed in Pi*Z liver tissue most mitochondria were in a swollen state with hypodense matrix (Figure 5B), consistent with our previous observation demonstrating AATD-associated hepatic mitochondrial dysfunction (24). Space of Disse was normal in Pi*M liver tissue (Figure 5C), while Pi*Z liver contained collagen fibers within the space of Disse (Figure 5D). As compared to Pi*M hepatocytes (Figure 5E), hepatocytes from the Pi*Z liver contained disorganized ER, while many bound ribosomes were still visible on the surface of the rough ER. Moreover, we observed increased interaction between ER and mitochondria, also known as mitochondria-associated membranes (MAMs) (25) (Figure 5F). Therefore, we analyzed levels of VDAC1 and GRP75 in liver homogenates of Pi*M and Pi*Z mice by western blotting (Figure 4G). These proteins are participating in the formation of MAMs. Semiquantitative analysis of western blotting confirmed GRP75 and VDAC1 levels were higher in Pi*Z compared to Pi*M liver tissue (GRP75: *p* = 0.0342; VDAC1: *p* = 0.0499) (Figure 5H). We also performed western blot analysis on the liver tissue lysates to determine changes in expression levels of ER chaperones and ER stress related genes in Pi*Z livers in comparison to Pi*M livers. Our analysis revealed significantly higher expression of BiP, CHOP, heat shock protein 60 (Hsp60), and ATF4 in the livers of Pi*Z as compared to Pi*M mice (Figure 5I).

**Figure 5.** Ultrastructure of the liver tissue from transgenic mice. (A) Transmission electron microscopy (TEM) images of a hepatocyte from Pi*M mouse liver tissue (n=1) and (B) Pi*Z mouse (n=1). LD: lipid droplet. (C) Space of Disse in Pi*M liver tissue as compared to (D) Pi*Z. Black arrow indicates collagen deposition in the space of Disse from Pi*Z mouse liver tissue. (E) Endoplasmic reticulum and mitochondrial morphology of the Pi*M as compared to (F) Pi*Z liver tissue. Black arrows indicate endoplasmic reticulum of the Pi*Z liver section. (G) The expression levels of VDAC1 and GRP75 in liver homogenates of Pi*M and Pi*Z mice as assessed by western blotting. (H) Quantification of the band intensities of VDAC1 and GRP75. (I) Western blot analysis on liver tissue lysates determining the expression levels of BiP, CHOP, heat shock protein 60 (Hsp60), and ATF4 in the liver of Pi*Z as compared to Pi*M mice (n=3). p-values have been calculated using student t-test method. Pi*M, normal variant of human α-1 -antitrypsin; Pi*Z, mutant human α-1 -antitrypsin.

### Single-nucleus RNAseq Characterization of Liver Tissue in the AATD Mouse Model

Liver tissue of Pi*M and Pi*Z mice were subjected to single-nucleus (Sn)-RNA sequencing (Figure 6A). According to the mouse cell marker expression profile curated in the SingleR package, 11 cell clusters were annotated and 5 main different liver cell types were recognized, including hepatocytes, endothelial cells, Kupffer cells, hepatic stellate cells, and colangiocytes (Figure 6B). Marker genes showed exclusively high expression in their corresponding cell types, such as Alb in hepatocytes, Clec4f in Kupffer cells, Rgs5 in hepatic stellate cells, Clec4g and Stab2 in sinusoidal endothelial cells, and Krt19 in cholangiocytes (Figure 6C). Hepatocytes had the largest population in captured cells (over 5,000 cells in both samples), while the second largest cell population was hepatic stellate cells (1,000 cells). We also observed higher numbers of cholangiocytes, Kupffer cells, and endothelial cells were found in the Pi*Z liver tissue and might be associated with liver injury caused by hepatic accumulation of α-1 antitrypsin (Figure 6D).

**Figure 6.** Single-nucleus characterization of the liver tissue from transgenic mice model. (A) Graphical representation of workflow for tissue sample collection and single nuclei RNA-sequencing. (B) Cell clusters generated by Seurat in R 4.3.2 and labeled with cell types identified using sc-type with R. (C) Distributions of genes marking each cell type, each shown on the same UMAP plot for comparison with the clusters in 5B. (D) Proportion of cells of each cell type found in each mouse with Pi*M found in blue and Pi*Z found in red. The black bars on the right show the number of cells of each cell type found in both mice combined. Figure has been generated by BioRender. Com.

### Transcriptional Profile of Liver Cell Types in AATD Associated Liver Disease

Hepatocytes, and hepatic stellate cells have been shown to play critical roles in chronic liver injuries, including AATD (6). Therefore, we compared transcriptional profiles of these cell types between Pi*M and Pi*Z livers. In the comparison between Pi*M and Pi*Z hepatocytes, 372 up-regulated and 622 down-regulated genes were identified (Figure 7A). These dysregulated genes were found to participate in cellular activation, immune response, activation of signaling pathways in response to cell surface receptors, and cellular proliferation, suggesting dysfunction in response to inflammatory stimuli in hepatocytes (Figure 7B). In comparing hepatic stellate cells from Pi*Z liver with Pi*M, 172 up-regulated and 105 down-regulated genes were identified (Figures 7C). These significantly altered genes were enriched in cellular differentiation, regulation of immune cells’ activation, and immune related processes, such as positive regulation of immune cells adhesion, and leukocyte migration (Figure 7D). We also compared Kupffer cells from the liver samples and found 36 up-regulated and 12 down-regulated genes in Pi*Z as compared to Pi*M liver tissue (Figure 8A). These significantly dysregulated genes were found to be mostly involved in lipid and fatty acid metabolism (Figure 8B). In comparing liver endothelial cells from Pi*Z liver with Pi*M liver tissue, 121 up-regulated and 99 down-regulated genes were identified (Figure 8C). These significantly altered genes were mainly related to metabolic changes and liver development (Figure 8D).

**Figure 7.** Transcriptional profile of hepatocytes and hepatic stellate cells from Pi*M and Pi*Z mice. (A) Heatmap showing up- and down-regulated genes for hepatocytes in Pi*M and Pi*Z samples obtained using Seurat version 5. (B) Gene ontology analyses of biological processes for hepatocytes generated using clusterProfiler. (C) Heatmap showing up- and down-regulated genes for hepatic stellate cells in Pi*M and Pi*Z samples obtained using Seurat version followed by (D) Gene ontology analyses of biological processes for hepatic stellate cells generated using clusterProfiler. Pi*M, normal variant of human α-1 -antitrypsin; Pi*Z, mutant human α-1 -antitrypsin.

**Figure 8.** Transcriptional profile of Kupffer cells and liver sinusoidal endothelial cells from Pi*M and Pi*Z mice. (A) Heatmap showing up- and down-regulated genes for Kupffer cells in Pi*M and Pi*Z samples obtained using Seurat version 5. (B) Gene ontology analyses of biological processes for Kupffer cells generated using clusterProfiler. (C) Heatmap showing up- and down-regulated genes for liver sinusoidal endothelial cells in Pi*M and Pi*Z samples obtained using Seurat version followed by (D) Gene ontology analyses of biological processes for liver sinusoidal endothelial cells generated using clusterProfiler. Pi*M, normal variant of human α-1 -antitrypsin; Pi*Z, mutant human α-1 -antitrypsin.

## DISCUSSION

AATD comprises a heterogeneous group of pathologic entities with the most common manifestations being lung inflammation and liver disease (26). Current mouse models of AATD have helped to progress understanding of pathologies and tissue injuries associated with AATD (27–30). However, these AATD mouse models express human α-1 antitrypsin protein at high levels in addition to endogenous expression of mouse α-1 antitrypsin (31). There is a need to improve AATD mouse models, including developing a model to study α-1 antitrypsin loss of function that better replicates AATD human disease (7). To overcome this limitation, we have generated a novel humanized transgenic model for AATD-associated liver disease on the background of a *SERPINA1*-null mouse (11). This model was established by microinjection of either the human M or Z variant of the *SERPINA1* gene into the oocytes of C57BL/6J-*SERPINA1* knockout mice. The resulting mice harboring the Z mutation of human α-1 antitrypsin gene display accumulation of α-1 antitrypsin polymers in the liver tissue and are susceptible to hepatic injury, including hepatotoxicity caused by ER stress and liver fibrogenesis, phenocopying key clinical features of AATD-associated liver disease. There is good agreement among species (i.e. mouse and human) regarding protein expression of α-1 antitrypsin in hepatocytes (32–34). Our humanized transgenic AATD mice are fully viable and express either the wildtype (M) or Z mutant variant of human α-1 antitrypsin in their hepatocytes (6). In AATD individuals carrying the Z allele, hepatic polymerization and aggregation of misfolded mutant α-1 antitrypsin correlates with hepatotoxicity and low plasma levels of α-1 antitrypsin (35). Here, we demonstrate that transgenic Pi*Z mice show accumulation of polymerized α-1 antitrypsin in the liver. We also observed that retention of misfolded α-1 antitrypsin in the liver results in low levels of plasma circulating human α-1 antitrypsin in Pi*Z mice as compared to Pi*M control mice. Thus, our new AATD mouse model mimics not only the liver accumulation of mutant α-1 antitrypsin, but also changes in the plasma levels of α-1 antitrypsin observed in AATD patients, recapitulating circulating human AAT turnover.

Alpha-1 antitrypsin exhibits anti-inflammatory properties independent of anti-protease activity, playing pivotal roles in the regulation of immune cell differentiation and activation, and tissue repair (36). Here by comparing the liver tissue from *SERPINA1*-null and Pi*Z mice with WT, we show a significant increase in the number of inflammatory monocytes in the liver of null mice. We also observed lower neutrophil and non-inflammatory monocyte recruitment in the liver of null mice at baseline and in response to LPS induced inflammation that was associated with increased LPS-induced hepatotoxicity in null mice as compared to WT, by an increase in plasma levels of AST and ALT. This suggests that lack of α-1 antitrypsin results in an inflammatory phenotype in the null livers. Previous studies have demonstrated a cooperation between neutrophils and macrophages that regulate resolution of inflammation and induce tissue repair (37). We show that absence of α-1 antitrypsin in the liver of null mice results in impaired initiation of inflammation as well as resolution which may affect liver repair after injury. We also investigated whether introducing human α-1 antitrypsin, either in its normal form or a mutated variant would alleviate or reverse the newly described inflammatory condition in the null liver. Our results revealed the normal but not the mutant human α-1 antitrypsin could mitigate this inflammation. The introduction of human α-1 antitrypsin as in Pi*M mice, the null liver’s inflammatory reversed to normal, homeostatic state of WT mice, suggesting that Alpha-1 antitrypsin replacement, as in augmentation therapy might be sufficient to correct the underlying inflammatory issues in the liver of these mice and AATD individuals. These findings do not rule out other factors or more complex mechanisms, like bone marrow leukocyte production and release, and hepatic endothelial - leukocyte adhesion and transmigration are likely involved in maintaining the inflammatory liver phenotype, beyond just the presence of α-1 antitrypsin. These analyses provide evidence that our new transgenic AATD mouse model - lacking endogenous α-1 antitrypsin - provides better tool to study AATD-associated liver disease by enabling to study both α-1 antitrypsin loss-of-function and toxic gain-of-function in the same animal model.

AATD-associated liver disease is predominantly characterized by liver fibrogenesis resulting from chronic hepatic injury (22, 38). The progression of liver fibrosis involves modulation of the liver extracellular space and changes of the composition and density of extracellular matrix (ECM) proteins (39), such as increased deposition of collagen and α-SMA (40). Collagen and α-SMA are markers for activation of hepatic stellate cells across multiple models of liver fibrosis (41), including AATD-associated liver disease (42). Consistent with human AATD-associated liver disease (42), IHC analyses revealed that the liver tissue from Pi*Z mice display significant deposition of collagen and α-SMA in the portal ECM, indicative of ECM remodeling in Pi*Z livers consistent with AATD and several other chronic liver diseases (43). Once again, these findings show compliance with the liver fibrogenesis, and phenotype required to a suitable animal model for AATD-associated liver disease.

Elevated plasma levels of cytokines represent a characteristic feature of chronic liver disease as a consequence of liver dysfunction (44). It has been shown that AATD is associated with mild systemic inflammation and elevated plasma levels of inflammatory mediators that may contribute to disease (42, 45, 46). The results presented in this study demonstrate that a significant number of cytokines/chemokines are elevated in the plasma of Pi*Z mice as compared to Pi*M. The higher concentrations of IL-18, IL-2, CCL2, and IL-23 in the plasma of Pi*Z mice suggests a potentially significant role of these cytokines in AATD mediated liver inflammation. These data also indicate that activated immune cells from different tissues, such as liver, can be the source of plasma inflammatory mediators in AATD-associated systemic inflammation. These cytokines are mostly initiators of acute phase inflammatory response in the liver, including apoptosis of hepatocytes and hepatic inflammation in chronic liver diseases (47, 48). IL-18, a pro-inflammatory cytokine, is known to cause liver toxicity (49, 50). It has been shown that IL-18 and IL-2 synergistically enhance proliferation and cytolytic activity of peripheral blood mononuclear cells (51). In addition, expression of CCL2 in alcoholic liver disease has been shown to be associated with liver disease severity (47). Altogether, these results suggest in Pi*Z mice elevated plasma levels of cytokines associate with progression of AATD-associated liver inflammation.

The hepatic ER plays a critical role in liver homeostasis, being the main synthesis site of proteins and lipids. Protein and lipid synthesis in the ER is required for appropriate function across cellular compartments, including mitochondria (52). We have previously shown that AATD mediated ER stress enhances formation and accumulation of lipid droplets associated with mitochondrial dysfunction in our previous AATD mouse model (24). Consistent with our previous report, our new Pi*Z transgenic mice show accumulation of lipid droplets in addition to α-1 antitrypsin globules within hepatocytes, which is associated with expression modifications of proteins involved in ER stress and MAMs formation. MAMs are the ER regions that mediate communication between the ER and mitochondria. They have a specific structure enriched in proteins and lipids involved in cellular homeostasis, survival, and metabolism (53). In Pi*Z animals, we found the number of MAMs in hepatocytes is increased compared to Pi*M. This suggests a compensatory mechanism in Pi*Z liver tissue aimed at enhancing molecule shuttling between the ER and the mitochondria in response to increased ER stress. Moreover, we have previously evidenced mitochondrial dysfunction associated with AATD liver disease, including enhanced energy-producing capacity in addition to changes of AATD-associated mitochondrial morphology (24). TEM analysis from our new transgenic animals also indicated dramatic changes in hepatic mitochondrial morphology of Pi*Z liver tissue. The shift in mitochondrial distribution to larger sizes and loss of complexity provide solid evidence of injured mitochondria. From morphological assessment, disorganization of cristae are found. The disruption in the organization of the mitochondria likely causes mitophagy, clearing and recycling damaged mitochondria from the cell (52).

Our study revealed substantial levels of ER stress in the Pi*Z mouse liver tissue due to hepatic accumulation of misfolded α-1 antitrypsin. α-1 antitrypsin exerts its effects mainly through the mechanism regulating autophagy (54) and unfolded protein response (55). α-1 antitrypsin is also known to bind several chaperones, including BiP, in the hepatocytes (56, 57). Consistent with previous reports, we observed that in the liver of Pi*Z mice, impaired folding and secretion of mutant α-1 antitrypsin results in sustained ER stress, leading to upregulation of ER chaperones involved in α-1 antitrypsin trafficking. Together with previous reports showing ablation of key molecules in ER stress response links impaired α-1 antitrypsin trafficking and polymerization, our data suggest that ER stress response is induced physiologically during the expression of Z mutant α-1 antitrypsin in the Pi*Z hepatocytes. Importantly, downstream molecules involved in ER stress responses may fail to be fully induced or activated in AATD-associated liver disease despite activation of upstream ER stress sensors, reflecting enhanced ER stress. On the other hand, when ER stress sensors are not fully activated for some reason, the downstream ER stress responses are not sufficiently induced, even under enhanced ER stress (58), known as ER stress response failure (59).

Single-cell RNA sequencing is a very powerful tool to assess gene expression at the fundamental level of an individual cell. SnRNA-seq is also a single cell sequencing approach that bypasses the enzymatic cell dissociation step to study complex tissues at single cell level (60, 61). Here, we performed single-nucleus transcriptomics of Pi*M and Pi*Z livers to capture parenchymal cells in high resolution and their interplay with the natural inactivated state of non-parenchymal cells (62) in liver injury associated with AATD. We observed slight loss of hepatocytes in addition to an increase in the proportion of Kupffer cells in the Pi*Z liver compared to Pi*M. The reduced number of hepatocytes may indicate liver fibrogenesis and impaired liver regeneration (63) caused by AATD-mediated hepatotoxicity. The increase in the proportion of inflammatory cells has been shown to promote progression of liver inflammation and fibrosis in chronic liver injury (64). Furthermore, our snRNA-seq transcriptional analysis revealed a difference of expression patterns between Pi*M and Pi*Z livers’ transcriptome and identified some gene markers of liver injury associated with AATD. We observed in Pi*Z liver, hepatocytes, hepatic stellate cells, liver sinusoidal endothelial cells, and Kupffer cells expressed genes related to proliferation, activation, and cell-cell adhesion. The expression of genes related to cell proliferation and cell cycle division is higher in Pi*Z than in Pi*M liver tissue, indicating the proliferation ability of hepatocytes is enhanced after hepatic injury to compensate for hepatocyte loss in the AATD liver. This might also be a signal for malignant transformation of Pi*Z liver tissue. These data suggest hepatotoxicity mediated by AATD promotes proliferation and activation of immune cells in the Pi*Z liver in response to injury.

During liver fibrogenesis, either liver-resident cells or patrolling and infiltrating leukocytes attracted to the site of liver damage, interact with each other mostly by direct cell–cell contact mediated by cell adhesion molecules. In addition, cell adhesion molecules support binding of cells to the ECM. Our data suggest overexpressed hepatic adhesion molecules in the Pi*Z liver may be involved in liver fibrogenesis by recruiting immune cells to the AATD liver. Our study suggests single-cell transcriptional alterations in the liver tissue of our AATD-associated liver disease mouse model. Although our findings are based on mouse models and need validation in clinical patient samples, these analyses may provide cognition of molecular pathological changes and potential therapeutic targets for AATD-associated liver disease.

## CONCLUSIONS

It should be appreciated that AATD transgenic mice were previously generated (6) in which the human α-1 antitrypsin gene was inserted in mice with wildtype background as a model for studying the role of α-1 antitrypsin in AATD-associated liver disease. However, in these mice there was no deletion of mouse endogenous α-1 antitrypsin. In our AATD mouse model, the purified DNA, containing the entire human *SERPINA1* gene, was injected into eggs from *SERPINA1* knockout mice. As a result, these new humanized AATD mice lack the endogenous mouse α-1 antitrypsin protein, avoiding potential interference of the endogenous gene. Results from the present study demonstrate that compared to Pi*M mice, our new Pi*Z model presents with AATD liver toxic gain-of-function and associated liver fibrogenesis as a result of hepatic accumulation of misfolded α-1 antitrypsin. Moreover, we describe a novel α-1 antitrypsin loss-of-function associated with a pro-inflammatory liver phenotype, ameliorated by introduction of normal, M, α-1 antitrypsin protein, as in Pi*M mice. These humanized AATD mice should prove a valuable model in future studies investigating the role, relevance, and regulation of the α-1 antitrypsin gene, the mechanisms underlying AATD-associated liver disease, and in the drug discovery process.

## STUDY LIMITATIONS

This study has several limitations. (A) This model is limited to 4-6 months of AATD-associated liver phenotype investigation. Future studies will be needed to determine if older mice will maintain the AATD-associated liver phenotype. (B) Although AATD is multi organ disease, we treated this model as a chronic liver disease model by only assessing liver tissue phenotype. Investigation of other organs, including lungs, will be needed to fully characterize this animal model. (C) We have reported abnormalities in the Pi*Z liver’s mitochondrial morphology. Mitochondrial electron transport chain and respiratory complex analysis warrants further study. (D) We obtained transcriptomic data from a single animal in each group. To validate our findings, not only more samples will be needed, but more time intervals during the pathological process will be necessary. (E) The histological features of the liver tissues and fibrosis phenotypes will require additional delineation of potentially unique mechanisms of fibrosis development or progression. (F) Comparison of male and female mice did not reveal differences in histopathological data, demonstrating that the AATD-induced liver phenotype in our transgenic mouse model is not dependent on sex. (G) Finally, this study did not focus on other treatment strategies than α-1 antitrypsin replacement, which we show it ameliorates liver inflammation.

## ACKNOWLEDGMENTS

The authors acknowledge the University of Florida’s ICBR core technician Rudy Alvarado (Electron Microscopy Core, RRID:SCR_019146) for his assistance in training and acquisition of microscopy images in this manuscript. They also thank the Singulomics Corporation for assistance in gene sequencing and gene expression analysis. The authors thank the University of Florida Molecular Pathology Core (RRID:SCR_016601) for sectioning and staining of liver tissues. BioRender has been used to generate schematic pictures.

---

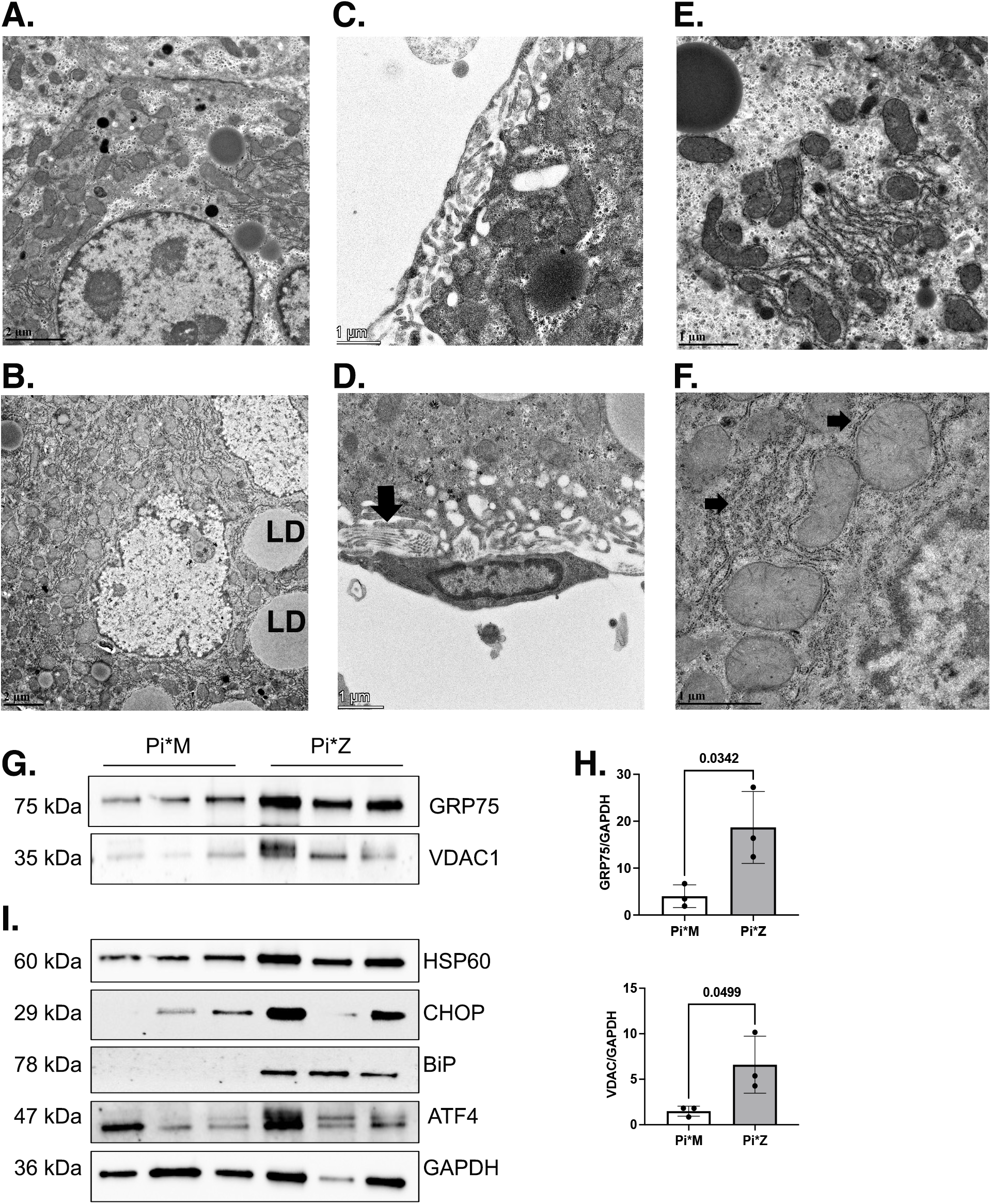

